# Abnormality Detection for Medical Images Using Self-Supervision and Negative Samples

**DOI:** 10.1101/2023.05.29.542748

**Authors:** Nima Rafiee, Rahil Gholamipoor, Markus Kollmann

**Author notes:** {, }. Equal contribution (Nima Rafiee, Rahil Gholamipoor).

## Abstract

Recent progress in computer-aided technologies has had a considerable impact on helping experts with a reliable and fast diagnosis of abnormal samples. In particular, self-supervised and self-distillation techniques have advanced automated out-of-distribution (OOD) detection in the image domain. Further improvements in OOD detection have been observed by including negative samples derived from shifting transformations of real images. In this work, we study different ways of creating negative samples for medical images and how effective they are when leveraging them in a self-supervised self-distillation framework. We investigate the impact of various types of negative examples by applying different shifting transformations on samples when they are derived from in-distribution training data, from an auxiliary dataset, or a combination of both. For the case of the auxiliary dataset, we compare the OOD detection performance when auxiliary samples are extracted from an in-domain or an out-domain. Our approach uses only data belonging to healthy people during the training procedure and does not require any additional information from labels. We demonstrate the efficiency of our technique by comparing abnormality detection performance on diverse medical datasets, setting new benchmarks for pneumonia, polyp, and glaucoma detection from X-ray, colonoscopy, and ophthalmology images.

## 1 Introduction

In recent years, computer-aided diagnosis in medical image screening has gained increased attention. In particular, detecting whether a sample includes some abnormality can help medical experts with faster and more reliable decisionmaking. Diagnosis problems can be frequently assigned to the problem of out-of-distribution (OOD) detection in machine learning and statistical inference. OOD detection or anomaly detection refers to the problem of detecting if a test sample has the same distribution as training data or is drawn from a different distribution. Diverse techniques developed for computer vision problems have been successfully applied to abnormality detection in the medical field. In [26, 30], deep supervised methods are used to classify X-ray and colonoscopy images. Despite the promising results, these approaches rely on annotated samples for abnormalities that are not available or only available to a very limited number. Typically, the number of healthy samples outnumbers abnormal ones, which results in a challenging unbalanced classification problem. To overcome these problems, many studies have investigated the use of unsupervised or semi-supervised methods [7, 27, 20]. These methods aim at detecting abnormalities by learning the distribution of healthy/normal data. A well-studied category of unsupervised methods named Variational Autoencoders (VAEs) [13] uses reconstruction error. The assumption is that abnormal samples can not be reconstructed equally accurately as training data (lower likelihood), where the model only uses normal images during the training. However, it has been shown that in practice, these models can be prone to reconstruct the abnormal samples fairly well, which lowers the detection performance [26, 2]. Furthermore, it is shown that these density estimation-based methods can assign a higher likelihood to OOD samples compared to in-distribution (in-dist) test data [19]. Recently the effectiveness of self-supervised learning has received considerable attention in different domains, such as the visual domain [5], which enables learning robust representations from unlabeled data. Due to their efficiency, self-supervised pre-text tasks such as predicting geometric transformations [10] or contrastive learning [28, 31, 9, 23, 25] have been designed for OOD detection in both natural and medical images. In [25], negative samples, drawn from shifting transformation of train data, are incorporated into a contrastive method to further tighten the decision boundary around normal samples resulting in an improved OOD detection score. This approach is also supported by Ref. [11], where supervised and density estimator models are exposed to some auxiliary datasets and negative samples. In [21], a self-distillation approach similar to DINO [4] is used with negative samples in order to compensate for the limitations of contrastive-based methods. Despite the numerous studies of leveraging negative samples in natural images, we believe it has remained untapped in the field of medical image processing. In this study, we investigate several techniques for generating negative examples for abnormality detection through the application of shifting transformations on in-dist train data, an auxiliary dataset, or both. We posit that these different negative samples result in an over-representation of specific features of in-distribution data and, accordingly, evaluate their effects on the performance of detecting abnormal samples using a self-supervised and self-distillation method. [21] demonstrated the effectiveness of rotation for natural object-centric images, as it modifies high-level semantics while retaining low-level statistics. However, we show that the efficacy of creating negative samples varies across different datasets. In the case of ophthalmology samples, for example, rotation may not be effective as high-level semantics may remain unchanged, while other transformations, such as splitting the image into multiple patches, can be more effective. Due to the limited availability of medical imaging datasets, we investigate the use of out-domain datasets such as ImageNet [**?**] for generating negative samples, comparing their efficacy to that of an in-domain auxiliary data source. Our methodology is applied to abnormality detection for three medical datasets, specifically the detection of pneumonia, polyps, and glaucoma from chest X-ray, colonoscopy, and ophthalmology images, respectively, with access only to normal samples.

## 2 Method

In this section, we describe our proposed approach, Fig. 1. Similar to [4], our framework uses teacher and student networks that have the same architecture, Vision Transformer [8] (ViT), and use distillation during training. Student and teacher are parameterized by two identical networks *g*_*s*_ = *g*(*x*|*θ*_*s*_) and *g*_*t*_ = *g*(*x*|*θ*_*t*_) which have different sets of parameters. For an augmented input image *x*, both student and teacher output *K*-dimensional vectors including soft-classes.

**Fig. 1:**
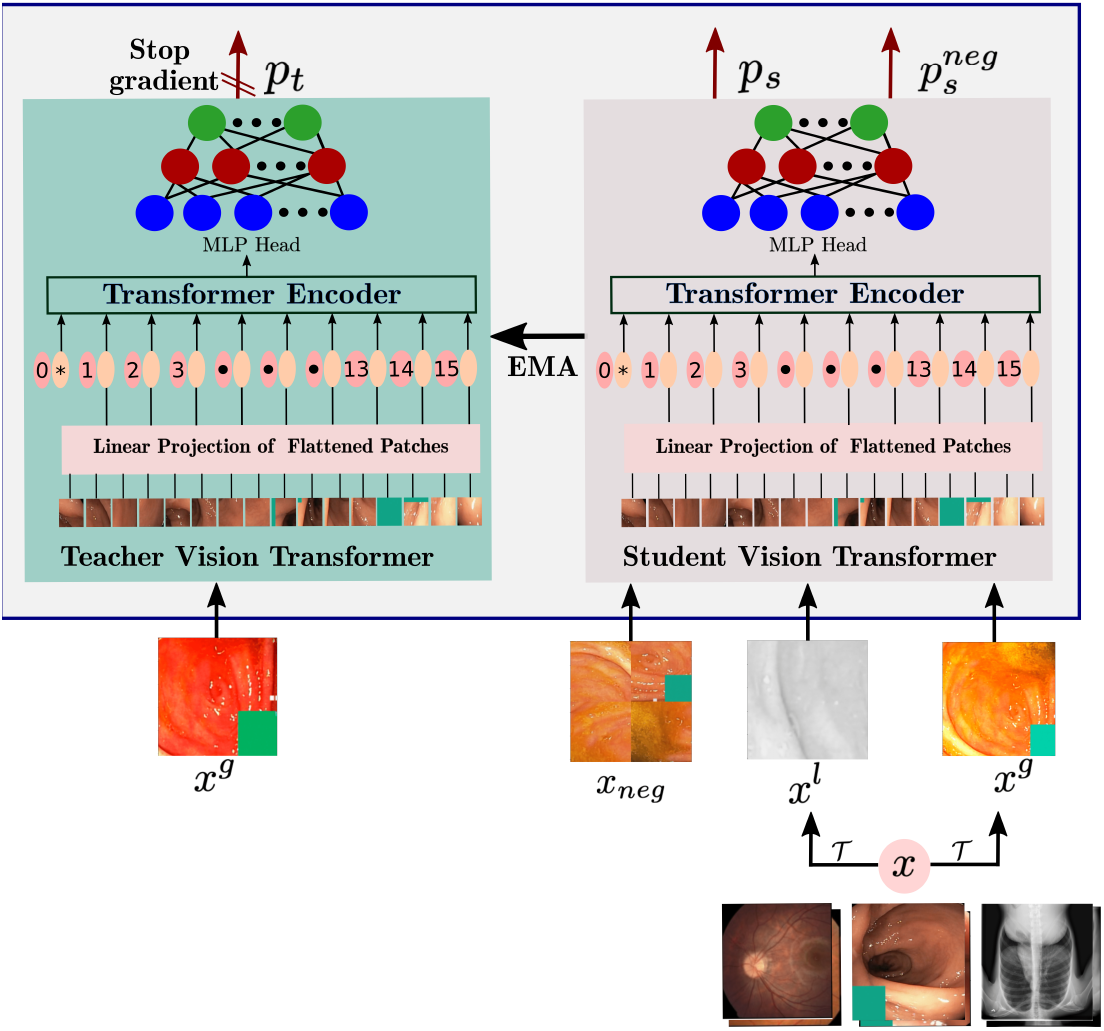
Overview of the proposed self-supervised framework, comprising student network (right) and teacher network (left). Student and teacher map two randomly augmented views of the same image to the same class. *x*^*g*^ and *x*^*l*^ are global and local views of image *x* where *x*^*g*^ *∼* 𝒯(*x*) and *x*^*l*^ *∼ 𝒯*(*x*). Negative sample, *x*_*neg*_, is generated by applying first a shifting transformation, such as random rotation, followed by *𝒯* to either an in-dist image *x* or an auxiliary image *x*_*aux*_.

The probability of *x* falling in soft-class *k* is computed using a temperature-scaled softmax function defined as

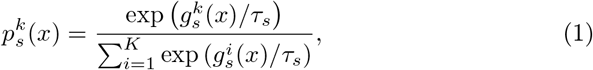

where *τ*_*s*_ *>* 0 is student temperature. The same formula, Eq. 1, holds for teacher with temperature *τ*_*t*_. The student parameters are updated by back-propagating the gradients through the student network, while the teacher parameters are updated with the Exponential Moving Average (EMA) of the student parameters.

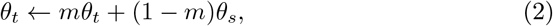

where 0 ≤ *m ≤* 1 is a momentum parameter. For *τ*_*t*_ *< τ*_*s*_, the training objective is given by the cross entropy (CE) loss for two non-identical transformations *x*^*′′*^, *x*^*′*^ of an image *x* drawn from the training set, *𝒟*_*train*_

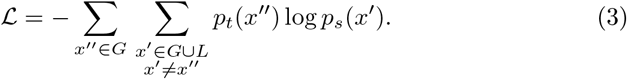

We additionally use the multi-crop strategy [3], wherein *M* global views 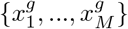 and *N* local views, 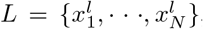, are generated based on a set of transformations 𝒯. Global views usually cover a larger region of the original image, while local views cover smaller as they are the results of stronger cropping. All global and local views are passed through the student network, while the teacher has only access to the global views encouraging local-to-global correspondence.

The CE loss, Eq. 3, is minimized such that two transformed views of an input image are assigned to the same soft-class. The applied transformations 𝒯 are chosen to be strong and diverse enough such that the generated images generalise well over the training data. The transformations are designed in order to learn higher-level features and semantic information and avoid learning lower-level features.

### 2.1 Auxiliary objective for OOD detection

For OOD detection, the representations should be enriched by in-dist specific features and deprived of features that frequently appear in other distributions from the same domain. This can be achieved by designing a negative distribution *𝒟*_*neg*_ that keeps most of the low-level features of the in-dist data but changes the high-level semantics.

In addition to the self-distillation objective, Eq. 3, we define an auxiliary task to encourage the student to have a uniform softmax response for negative examples. This can be done by a similar objective as Eq. 3 when temperature *τ*_*t*_ *→ ∞*

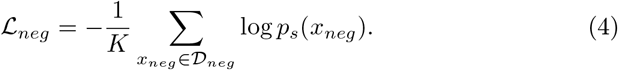

The total loss of our proposed method is defined by a linear combination of the two objectives

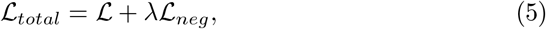

where *λ >* 0 is a balancing hyperparameter.

### 2.2 Negative samples

A negative distribution *𝒟*_*neg*_ can be realised by additionally applying shifting transformations to samples from *𝒟*_*train*_ or from an auxiliary set augmented by *𝒯*. We consider the following shifting transformations to shape *𝒟*_*neg*_.

– NoNeg: no negative samples are included (*λ* = 0).

– Rot: samples are randomly rotated by *r∼ 𝒰* ({90^*°*^, 180^*°*^, 270^*°*^}).

– Rot-360: rotation by an angle randomly sampled from range (0^*°*^, 360^*°*^).

– Perm-*n*: random permutation of image patches where the image is sliced in *n* square patches.

– Pixel-Shuffle: randomly shuffles all pixels in the image.

### 2.3 Evaluation protocol for OOD detection

Different metrics have been used in various studies for detecting Out-of-Distribution (OOD) samples, with Mahalanobis distance and cosine similarity being among the most commonly used ones [25, 23, 9]. In [9], it is shown that applying Mahalanobis distance on the self-supervised learned representation achieves promising results for the problem of pneumonia detection on X-ray images. For natural object-centric colorful images, both cosine similarity and Mahalanobis distance have been found to be effective for OOD detection problems [21, 23, 25]. In this work, for the problem of pneumonia detection using X-ray images, we use Mahalanobis distance to ensure a fair comparison with the contrastive self-supervised method used in [9], and for two other datasets with colorful images, we use cosine similarity score. Additionally, we compare the results of these two metrics for all three datasets in section 4. To calculate scores, we drop the fully connected head and use normalised ViT backbone output as feature vector *f* for calculating evaluation scores. For each given test sample *x*, we calculate Mahalanobis distance-based anomaly score, *S*_*md*_(*x*), as

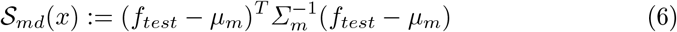

where *μ*_*m*_ and *Σ*_*m*_ are the mean and covariance of the all feature vectors *f*_*m*_ from the training data, *𝒟*_*train*_. We calculate the cosine similarity based anomaly score *𝒮*_*cs*_(*x*) for test sample *x*

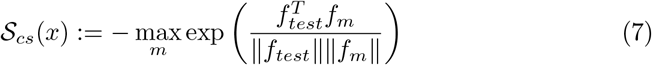

Detection is assessed with Area Under the Receiver Operating Characteristic curve (AUROC).

## 3 Experimental Setup

### 3.1 Dataset

We assess our model performance on three different health screening medical imaging benchmarks, chest X-ray images, colonoscopy images, and fundus images for glaucoma detection.

#### RSNA

The Radiological Society of North America (RSNA) Pneumonia Detection Challenge dataset [24] is a publicly available dataset of frontal view chest radiographs. Each image was labeled as “Normal”, “No Opacity/Not Normal” or “Opacity”. The Opacity group consists of images with opacities suspicious for pneumonia, and images labeled “No Opacity/Not Normal” may have lung opacity but no opacity suspicious for pneumonia. The RSNA dataset contains 26, 684 X-rays with 8, 851 normal, 11, 821 no lung opacity/not normal and 6, 012 lung opacity.

#### Hyper-Kvasir

The Hyper-Kvasir dataset is the largest public gastrointestinal dataset [1]. The data were collected during real examinations and partially labeled by experienced endoscopists. The dataset contains 110, 079 images from patients, with 10, 662 labeled images. Following [28], we take 2, 100 images from “cecum”, “ileum”, and “bbps-2–3” cases as normal and 1000 abnormal images from “polyp” as abnormal. We take 1, 600 images for the training set and 500 images for the test set.

#### LAG

The LAG dataset is a large-scale image dataset for glaucoma detection [15], containing 4, 854 images with 1, 711 positive glaucoma (abnormal) and 3, 143 negative glaucoma (normal) scans. For consistent comparison, following [28], we take 2, 343 normal images for training and 800 normal images and 1, 711 abnormal images for testing.

### 3.2 Auxiliary Dataset

For the auxiliary dataset, we compare the use of samples from ImageNet as an out-domain dataset or from an in-domain one if any are available. For the RSNA dataset of X-ray images we use CheXpert [12], and for the Hyper-Kvasir dataset of colonoscopy images all unlabeled Hyper-Kvasir images are taken as in-domain. For the LAG dataset, we only use ImageNet due to the unavailability of any in-domain dataset. We highlight that we do not use any label information to shape negative samples.

### 3.3 Training

Our proposed method has the same structure as DINO implementation. we use ViT-Small (ViT-S) backbone for all different training data. A patch size of 16 and *N* = 8 local views for both positives and negatives, but two global positive views and one global negative view are used. All global views are resized to 256×256 while local views to 96×96. The temperatures for teacher and student network are set to *τ*_*t*_ = 0.01 and *τ*_*s*_ = 0.1. During training, *τ*_*t*_ is linearly decreased from 0.04 to 0.01 in each epoch. *λ* and *K* are set to 1 and 4096 respectively for all our experiments. We use the Adamw optimizer [16] with an effective batch size of 256. For the base learning rate lr_base_, we use 0.001 for Hyper-Kvasir and LAG datasets and 0.002 for RSNA. For each dataset, we trained the model for 700 epochs. We conducted our experiments using 4 NVIDIA-A100 GPUs with 40 GB of memory. The image augmentation pipeline *𝒯* is based on DINO, except that for Hyper-Kvasir dataset, we rotate all positive views with the same randomly chosen angle to avoid information leaking from the position of existing green boxes in images. Finally, weight decay and learning rate are scaled with a cosine scheduler.

## 4 Experimental Results

### 4.1 Pneumonia Detection

For the problem of detecting pneumonia samples, we compare our proposed method with unsupervised methods trained on only healthy images. In line with [9], Mahalanobis distance *𝒮*_*md*_ is used as the anomaly score. As shown in Table 1, on the RSNA dataset, our method outperforms the previous unsupervised and self-supervised methods. we report results for the cases where OOD samples belong to No Opacity/Not Normal (“No Opacity” in Table 1) and Opacity. In Table 2, we inspect the anomaly detection performance on the Hyper-Kvasir dataset for polyp detection and on the LAG dataset for glaucoma detection. Our method can surpass the recently proposed self-supervised anomaly detection method, CCD [28], on both polyp and glaucoma detection, where we take *S*_*cs*_ as anomaly score.

**Table 1:** AUROC of OOD detection method trained on **RSNA** dataset

| Method | $\mathcal{D}_{ood} : \text{Opacity}$ | $\mathcal{D}_{ood} : \text{No Opacity}$ |
| --- | --- | --- |
| *Unsupervised methods trained on normal samples* |  |  |
| UAE [17] | 0.89 | 0.78 |
| Deep AD [18] | 0.838 | 0.704 |
| [9] | 0.940 | 0.828 |
| **Ours** | **0.941** | **0.841** |

**Table 2:**
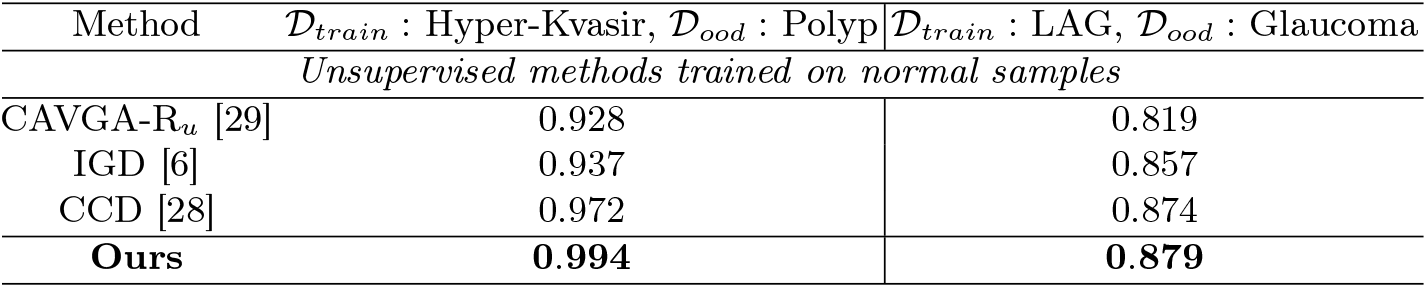
AUROC results on **Hyper-Kvasir** and **LAG** datasets

| Method | $\mathcal{D}_{train}$ : Hyper-Kvasir, $\mathcal{D}_{ood}$ : Polyp | $\mathcal{D}_{train}$ : LAG, $\mathcal{D}_{ood}$ : Glaucoma |
| --- | --- | --- |
| *Unsupervised methods trained on normal samples* |  |  |
| CAVGA-$R_u$ [29] | 0.928 | 0.819 |
| IGD [6] | 0.937 | 0.857 |
| CCD [28] | 0.972 | 0.874 |
| **Ours** | **0.994** | **0.879** |

### 4.2 Polyp and glaucoma detection

In Table 2, we inspect the anomaly detection performance on the Hyper-Kvasir dataset for polyp detection and on the LAG dataset for glaucoma detection. Our method can surpass the recently proposed self-supervised anomaly detection method, CCD [28], on both polyp and glaucoma detection, where we take *S*_*cs*_ as anomaly score.

### 4.3 Different shifting transformations

In Table 3, the impact of different shifting transformations, as explained in section 2.2 is explored. We found out that for RSNA and LAG datasets shifting transformations such as Rot achieve better performance than excluding negative samples or using other transformations such as Pixel-Shuffle. Our findings for these two datasets are in line with results reported in [21] where rotation as shifting transformation achieves the best performance for datasets of natural images such as CIFAR10 [14]. For the Hyper-Kvasir dataset, we found that applying Perm-4 as shifting transformation results in a better performance compared to Rot. Although, using Rot still achieves superior results compared to excluding the negative examples from training. Note that images in the Hyper-Kvasir dataset, have a green square box in the bottom right corner thus applying rotation on negative examples can result in an information leak and provide a shortcut for the model to minimize the loss. To avoid this information leak, positive views are rotated by the same angle randomly selected from *𝒰*({90^*°*^, 180^*°*^, 270^*°*^}) when we use Rot as a shifting transformation and we skip applying Rot-360 as a shifting transformation.

**Table 3:** The impact of different shifting transformations on AUROC results. Reported scores are for *𝒮*_*md*_ (*𝒮*_*cs*_).

| In-dist Dataset | NoNeg | Shifting transformations |  |  |  |  |
| --- | --- | --- | --- | --- | --- | --- |
|  |  | Rot | Rot-360 | Perm-4 | Perm-16 | Pixel-Shuffle |
| RSNA | 0.925(0.888) | **0.941(0.764)** | 0.933(0.766) | 0.924(0.634) | 0.908(0.887) | 0.925(0.733) |
| LAG | 0.799(0.862) | **0.849(0.879)** | 0.831(0.866) | 0.807(0.873) | 0.814(0.881) | 0.797(0.860) |
| Hyper-Kvasir | 0.974(0.875) | 0.989(0.915) | — | **0.996(0.994)** | 0.985(0.960) | 0.994(0.985) |

### 4.4 Different sources of negative samples

In Table 4, we examine the effect of creating negative samples by applying shifting transformations on samples from in-dist training, auxiliary dataset, or a combination of both. For the RSNA and LAG datasets, as it is shown, the AUROC score increases where a combination of both is used, while for Hyper-Kvasir, we see no difference. Moreover, the use of only auxiliary datasets shows slightly better performance for RSNA compared to only taking in-dist negative samples. On the other hand, for the LAG dataset, using in-dist negative samples results in better performance. The reason can be that for RSNA, the in-domain auxiliary datasets are from a broader distribution compared to in-dist train data with a higher chance of resembling OOD samples, but for the LAG dataset, even though the ImageNet dataset has a broader distribution, in-dist negatives are harder negative samples which can be more advantageous [22].

**Table 4:** AUROC results based on *𝒮*_*md*_ score for different negative sets where generated from in-dist train data, an auxiliary dataset or a combination of both.

| Train data | Negative sets |  |  |
| --- | --- | --- | --- |
|  | In-dist | Auxiliary | Combined |
| RSNA | 0.935 | 0.937 | 0.941 |
| LAG | 0.84 | 0.832 | 0.849 |
| Hyper-Kvasir | 0.996 | 0.996 | 0.996 |

### 4.5 In-domain and Out-domain auxiliary datasets

The evaluation of taking in-domain or out-domain auxiliary datasets is shown in Table 5. For RSNA X-ray images, OOD detection performance is improved by a large margin when negative samples are from an in-domain auxiliary set. However, for Hyper-Kvasir, the out-domain auxiliary has approximately the same performance as the in-domain.

**Table 5:** AUROC results across different auxiliary datasets where we take images from an in-domain medical dataset or ImageNet dataset as an out-domain.

| Train data | Negative sets |  |
| --- | --- | --- |
|  | ImageNet | Domain-related |
| RSNA | 0.912 | 0.941 |
| Hyper-Kvasir | 0.993 | 0.996 |

## 5 Conclusion

In this study, we present a self-supervised method that leverages self-distillation and negative samples for the task of abnormality detection without accessing label information. We study different ways of creating negative samples by applying shifting transformations on in-dist training data, an auxiliary dataset, or a combination of both. Additionally, we compare the impact of having auxiliary samples from domain-related distribution or from a different domain such as ImageNet. Moreover, we compare the abnormality detection performance using two different evaluation metrics including cosine similarity and Mahalanobis distance. A major motivation behind this work is that we take only normal samples during training which makes our method suitable for yet unknown abnormalities. In anomaly detection, our method outperforms SOTA methods on the RSNA, Hyper-Kavsir, and LAG datasets.

